# Predicting the Influence of Axon Myelination on Sound Localization Precision Using a Spiking Neural Network Model of Auditory Brainstem

**DOI:** 10.1101/2021.10.15.464581

**Authors:** Ben-Zheng Li, Sio Hang Pun, Mang I Vai, Tim C. Lei, Achim Klug

**Affiliations:** Department of Physiology and Biophysics, University of Colorado Anschutz Medical Campus, Aurora, CO 80045 USA; Department of Electrical Engineering, University of Colorado Denver, Denver, CO 80204 USA; State Key Laboratory of Analog and Mixed Signal VLSI, University of Macau, Macau; Department of Electrical and Computer Engineering, Faculty of Science, University of Macau, Macau

**Keywords:** spatial hearing, spiking neural network, sound localization, myelination, medial superior olive, auditory brainstem

## Abstract

Spatial hearing allows animals to rapidly detect and localize auditory events in the surrounding environment. The auditory brainstem plays a central role in processing and extracting binaural spatial cues through microsecond-precise binaural integration, especially for detecting interaural time differences (ITDs) of low-frequency sounds at the medial superior olive (MSO). A series of mechanisms exist in the underlying neural circuits for preserving accurate action potential timing across multiple fibers, synapses and nuclei along this pathway. One of these is the myelination of afferent fibers that ensures reliable and temporally precise action potential propagation in the axon. There are several reports of fine-tuned myelination patterns in the MSO circuit, but how specifically myelination influences the precision of sound localization remains incompletely understood. Here we present a spiking neural network model of the auditory brainstem with myelinated axons to investigate whether different axon myelination thicknesses alter the sound localization process. Our model demonstrates that axon myelin thickness along the contralateral pathways can substantially modulate ITD detection. Furthermore, optimal ITD sensitivity is reached when the MSO receives contralateral inhibition via thicker myelinated axons compared to contralateral excitation, a result that is consistent with previously reported experimental observations. Our results suggest specific roles of axon myelination for extracting temporal dynamics in ITD perception, especially in the pathway of the contralateral inhibition.

## 1 Introduction

In the mammalian brain, the precise temporal information encoded in action potentials and trains of action potentials is one major mechanism contributing to accurate neural integration and information processing. Such temporal precision is especially crucial for the localization process of sound sources in the auditory brainstem, especially for low-frequency sound sources. This process works through the microsecond-precise integration of signals between the two ears such that whenever the temporal precision is even slightly altered, sound localization accuracy suffers significantly (reviewed in Grothe, 2003; et al, Tollin & Yin, 2009; Grothe et al., 2010. Myelination of afferent fibers is a key mechanism in mammalian brains to ensure reliable, energy-efficient and temporally precise action potential propagation, and therefore, myelination is tightly controlled and actively managed in the brain (reviewed in Debanne, 2004). It is therefore not unexpected that such mechanisms have been described in the sound localization pathway as well (Stange-Marten et al. 2017; Seidl and Rubel, 2016; Ford et al., 2015).

In the auditory brain stem, sound localization is accomplished by the two principal localization nuclei, the lateral and the medial superior olive (LSO and MSO, respectively). High-frequency sounds are localized in the LSO by calculating the interaural intensity difference (ILD; Boudreau and Tsuchitani, 1966), while low-frequency sounds are localized in the MSO by calculating the interaural time difference (ITD; Goldberg and Brown, 1969). Mammals, including human listeners, are typically capable of resolving two sound sources that are just a few degrees separated from each other and, therefore, are capable of resolving ITD as small as several microseconds (Grothe, 2010). This extraordinarily high level of temporal precision is accomplished by a number of neural mechanisms, some of which ensure the propagation of action potentials with high temporal precision (Stange-Marten et al. 2017; Seidl and Rubel, 2016). Along the ITD pathway, action potentials generated in the cochleae of the two ears propagate in separate monaural pathways and eventually converge onto binaural neurons in the MSO, which is located several synapses upstream. How is this microsecond-level temporal precision preserved across multiple fibers, synapses, and several neuronal subtypes, and while one of the two inputs has a significantly larger distance to cover?

To accomplish temporally precise integration of binaural signals at the level of the MSO, several neural mechanisms along the afferent pathways have been described, including tuning of cochlear, synaptic, post-synaptic, and transmission delays (reviewed in Trussell, 1999; Grothe, 2003; et al, Tollin & Yin, 2009; Grothe et al., 2010). Notably, some experimental studies observed that the contralateral excitatory and inhibitory inputs to the MSO show different axon myelination patterns. The contralateral inhibitory pathway consists of axons with thicker layers of myelin, resulting in higher conduction velocities, compared to those of the ipsilateral excitatory pathway. These experimental results suggest that axonal myelination may be specifically adapted for tuning the input timing to the MSO, thereby actively contributing to spatial hearing perception (Schwartz, 1992; Stange-Marten et al. 2017; Seidl and Rubel, 2016; Ford et al., 2015).

While the structural fine tuning of myelination has been shown to control the conduction velocity of action potentials propagating between neurons, both physiologically and anatomically, axon myelination as a factor in circuit modeling is underexplored and is simply included as a constant in most models - probably due to inadequate understanding of the structure-function relationships (Nave et al., 2010). In this work, the role of myelin morphology as a contributing factor to the MSO sound localization circuit is specifically explored.

Prior modeling efforts of ITD coding at MSO primarily focused on the effects of post-synaptic integration of MSO neurons (Brand et al., 2002; Zhou et al., 2005; Leibold, 2010; Brughera et al., 2013; Myoga et al., 2014). Other studies modeled the axonal propagation time as a constant delay, not including axonal morphology (Wang et al., 2013; Encke and Hemmert, 2018). On the other hand, some studies included action potential timing difference by varying axonal propagation delays (Glackin et al., 2010; Pan et al., 2020). These models, however, were based on the Jeffress model of a delay line structure (Jeffress, 1948), which is anatomically inconsistent with neural inhibition observed in mammalian ITD circuits (Brand et al., 2002; Franken et al., 2015; Grothe & Pecka, 2014).

This study employed a spiking neural network (SNN) model to investigate how axonal structure and synaptic adaptation between the excitatory and inhibitory inputs to MSO can affect ITD perception. Specifically, the myelination thickness and the inhibitory conductance were modeled in detail and compared with pure tone and natural sound stimulation regarding ITD decoding accuracy and minimum temporal discrimination. Based on this SNN model and our decoding analysis, we found that the axon myelination patterns of both contralateral excitatory and inhibitory pathways can significantly modulate ITD decoding. The variation of myelin thickness, which results in conduction velocity variations along the excitatory pathways, can significantly shift the ITD tuning curve. On the other hand, axonal myelination and synaptic strength variations on the inhibitory pathway can significantly influence ITD sensitivity and precision.

## 2 Methods

### 2.1 Neuron and Synapse Model

The spiking neurons were modeled under a conductance-based leaky integrate-and-fire scheme. The membrane potential (*v*_*m*_) of a spiking neuron was described by the following first-order differential equation:

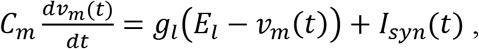

where *I*_*syn*_ is the total synaptic current comprising an excitatory and an inhibitory component:

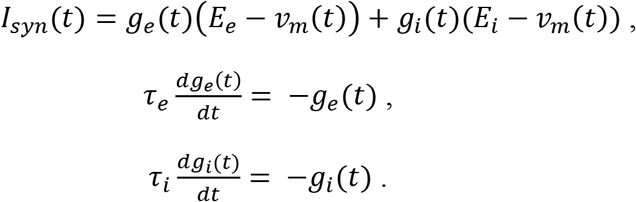

Descriptions of the parameters and the values used in the simulation are listed in Table 1. The excitatory and inhibitory synaptic conductances were modeled as first-order time decaying parameters with lifetimes of *τ*_*e*_ and *τ*_*i*_. When an action potential arrived at the presynaptic membrane, the conductance increased by Δ*g*_*e*_ or Δ*g*_*i*_ depending on the cell type, and subsequently decayed as described by the time constants. The spiking neuron will elicit an action potential when the membrane potential *v*_*m*_(*t*) reaches the firing threshold *v*_*th*_, and the membrane potential *v*_*m*_(*t*) is then reset to the resting potential *v*_*reset*_ after a short refractory period *τ*_*ref*_.

**Table 1.** List of Parameters for Neurons and Synapses

| Neuron parameter | Value | Description | Synapse Parameter | Value | Description |
| --- | --- | --- | --- | --- | --- |
| $C_m$ | 70 pF | membrane capacitance | $E_e$ | 0 mV | excitatory reversal potential |
| $V_{th}$ | -50 mV | threshold potential | $E_i$ | -70 mV | inhibitory reversal potential |
| $V_{reset}$ | -55.8 mV | reset potential | $\Delta g_e$ | 15 nS | excitatory postsynaptic conductance increment |
| $\tau_{ref}$ | 5 ms | refractory period | $\Delta g_i$ | 75 nS | inhibitory postsynaptic conductance increment |
| $E_l$ | -55.9 mV | leaky reversal potential | $\tau_e$ | 0.23 ms | excitatory time constant |
| $g_l$ | 13 nS | leaky conductance | $\tau_i$ | 2 ms | Inhibitory time constant |

### 2.2 Network Architecture

The architecture of the proposed SNN sound localization model (Figure 1A) consists of a left and a right MSO and related upstream nuclei. The cochlea first encodes acoustic stimuli, which are then sent as action potentials into the model through the auditory nerve (AN) and received by the Cochlear Nuclei (CN). At the CN, the Spherical Bushy Cells (SBCs) innervate the two MSOs bilaterally, and Globular Bushy Cells (GBCs) innervate the contralateral Medial Nucleus of the Trapezoid Body (MNTB) as well as the ipsilateral Lateral Nucleus of the Trapezoid Body (LNTB). Under this architecture, MSO cells receive bilateral excitation from the SBCs, ipsilateral inhibition from the LNTB and contralateral inhibition from the MNTB.

**Figure 1.**
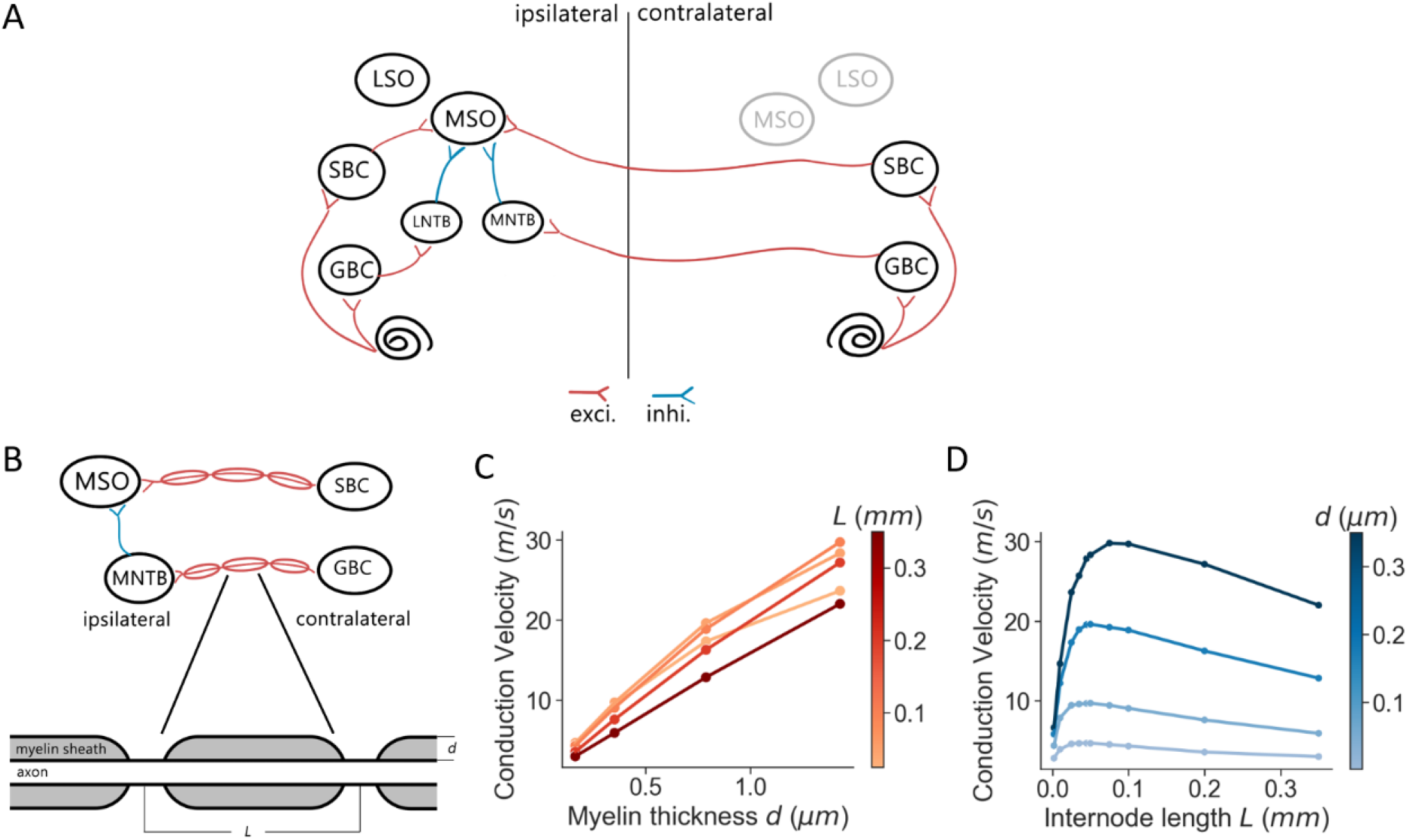
MSO model and myelination patterns. (A) Model architecture. (B) Myelination patterns along the contralateral pathway to MSO. (C, D) Axonal conduction velocity versus myelin thickness (C) and internode length (D).

Each neural population in the model contained 1000 spiking neurons that stochastically connected to neurons in other populations based on a connection probability (*p*_connect_). The synaptic connections were also associated with an axonal transmission delay (*t*_trans_) and a synaptic delay (*t*_syn_). These delays were normally distributed with a standard deviation of 0.05 ms. The transmission delays of the connections that traveled across the midline were derived from the corresponding conduction velocities and myelin thicknesses. The SNN model was implemented using the Brian2 simulator (Stimberg et al., 2019) and was simulated on a supercomputing cluster (RMACC Summit, University of Colorado Boulder). The simulation was repeated twenty times with different random seeds to compensate for random effects induced during stimuli encoding, network building, and decoding. Averaged metrics from all random permutations are reported as results. The specific parameters of the connections were adopted from previous studies (Brand et al., 2002; Spirou et al., 2005; Couchman et al., 2010; Roberts et al. 2013; Roberts et al. 2014; Encke and Hemmert, 2018) and are listed in Table 2.

**Table 2.** List of Parameters for Neurons and Synapses

| Population | Type | Connection | $p_{\text{connect}}$ | $\bar{t}_{\text{trans}}$ (ms) | $\bar{t}_{\text{syn}}$ (ms) |
| --- | --- | --- | --- | --- | --- |
| AN | Spike generator | AN → ipsi. GBC | 4 % | 0.2 | 0.6 |
| | | AN → ipsi. SBC | 4 % | 0.2 | 0.6 |
| SBC | Excitatory | SBC → ipsi. MSO | 0.6 % | 0.6 | 0.6 |
| | | SBC → cont. MSO | 0.6 % | *various*<sup>b</sup> | 0.6 |
| GBC | Excitatory | GBC → ipsi. LNTB | 0.3 % | 0.3 | 0.3 |
| | | GBC → cont. MNTB<sup>a</sup> | one-to-one | *various*<sup>c</sup> | 0.2 |
| MNTB | Inhibitory | MNTB → ipsi. MSO | 0.3 % | 0.1 | 0.3 |
| LNTB | Inhibitory | LNTB → ipsi. MSO | 0.3 % | 0.1 | 0.3 |
| MSO | Excitatory |  |  |  |  |

a: the Calyx of Held with $\Delta g_{\text{inhi}} = 250$ nS482 b: depends on the SBC myelin thickness $d_{\text{SBC}}$ 483 c: depends on the GBC myelin thickness $d_{\text{GBC}}$

### 2.3 Stimulus Encoding

For sound stimulation to the cochlea, two spike generators were used at the left and right AN to encode the sound signals into cochlear action potential responses. For pure tone stimulation, sinusoidal sound waves of 300 Hz with a 50 dB sound pressure level were generated with a sampling rate of 100 kHz. The duration of the sinusoidal wave was 100 ms with 20 ms ramp-up and ramp-down periods. For natural sound stimulation, sound samples were created based on 60 bird song clips of the long-eared owl collected from the Xeno-canto project (www.xeno-canto.org). The bird song clips were adjusted to 50 dB SPL with 20 ms ramp-up and ramp-down, and were up-sampled from 44.1kHz to 100kHz. Much of the previously published physiological data which informed our model were recorded in rodents such as mice or Mongolian gerbils. Thus, vocalizations of an owl species - a predator of these animal models - seemed appropriate.

Acoustic stimuli were sent to two ears with ITDs ranging from −1 to +1 ms with step sizes in log-scale. ITD is defined as the onset time difference of the same sound between the left and the right ear, with positive ITDs defined as sound leading at the right ear. For each ITD, the stimulation was repeated ten times for pure tones and eight times for natural sounds. A peripheral hearing model (Zilany et al., 2014; Rudnicki et al., 2015) was used to generate action potentials from the sound waves, and three spontaneous firing rates (high : medium : low = 6:2:2) were used to simulate action potential generation for AN composited auditory nerve fibers.

### 2.4 Axon Myelination and Conduction Velocity

The conduction velocity of myelinated axons was computed using the multi-compartment axon model of Halter and Clark (Halter and Clark, 1991; Ford et al., 2015; Arancibia-Cárcamo et al. 2017). In the following simulations, the axon internodal length was fixed to be 0.187 mm, and the node diameter was defined as 60% of the axon diameter. Meanwhile, the myelin thickness of the axon varied from 0.2 to 0.6 μm. The axon length from the SBC in CN to MSO on the ipsilateral side and from MNTB to MSO for adult gerbils were assumed to have a length of 4.5 mm estimated from the brain atlas of the adult Mongolian gerbil (Radtke-Schuller et al., 2016). As previously reported (Ford et al., 2015), the internodal length decreased at a distance of more than 0.5 mm from the branching area in MSO and 0.7 mm from the heminode near MNTB. At that point, the conduction velocity became more uniform along the rest of the axon, and the transmission delay was directly computed from the conduction velocity and the corresponding axon length with the uniform internodal length, i.e., 4 mm for the SBC-MSO projections and 3.8 mm for the GBC-MNTB projections.

### 2.5 Data Analysis

ITD responses to various sound stimuli were quantified as MSO firing rates in response to these stimuli. ITD tuning curves were computed by the trimmed mean from the 20% to 80% firing rates within the MSO population. For a more accurate peak firing rate analysis, the ITD tuning curves were interpolated with a precision of 0.1 μs and subsequently smoothed by a Savitzky–Golay filter with an 80 μs smoothing window. After the ITD curves were smoothed, the peak firing rate positions were then used to define the best ITD, the peak amplitude, and the Full-Width-at-Half-Maximum (FWHM) for the first few figures shown in the Results section.

For the ITD decoding analysis, the spike counts of MSO neurons during the course of the stimuli were used as decoding features. ITDs were predicted using a Support Vector Machine (SVM) classifier with a linear kernel trained with the leave-one-out cross-validation approach. Three hundred MSO neurons were then randomly selected from both left and right MSO and sent to the SVM classifier to predict ITD in response to each stimulation. The decoding accuracy and the mean squared error (MSE) were determined from the predicted ITDs with 17 ITD sub-classes in the range from –1 to 1 ms. The classification accuracy between 10 μs and –10 μs ITDs was also computed in the same manner to estimate the precision of the ITD detection.

The sensitivity of ITDs was accessed using the Just-Noticeable-Difference (JND) that quantifies the smallest perceptible change. It was computed by comparing ITD responses symmetrical to the zero time, e.g., –50 μs and 50 μs ITDs. For each pair of symmetrical ITD responses, the difference of firing rates between left and right MSO were compared using the one-tailed Mann-Whitney U test. The smallest symmetrical ITD that reached the minimum significant level indicates the JND.

## 3 Results

### 3.1 Conduction velocity varies with axon myelination patterns

Our overall results indicate that both the myelin thickness and the internodal length affect conduction velocity. While the level of myelin is directly proportional to the conduction velocity (Figure 1C), the internodal length is only correlated with the conduction velocity within a specific range (Figure 1D). Theoretically, increasing the myelin thickness should increase the axial current flow, which in turn increases the propagation speed of action potentials. On the other hand, when the internodal length is short, the increased internodal length results in a greater myelin coverage on the axon, which also increases the conduction velocity. When the internodal length becomes longer than a threshold length, the conduction velocity decreases with longer internodal length since the transfer efficiency of the depolarization between nodes is lower (Ford et al., 2015; Brill and Waxman, 1977). For the following results, the simulated axons had a constant internodal length as described in the Methods section, but the myelin thickness was varied for comparison.

### 3.2 SBC axon myelin thickness and ITD tunning

The firing rates of MSO neurons were studied during sinusoidal sound wave stimulation with varying ITDs, and were additionally recorded against changing myelin thicknesses of the SBC axon (*d*_*SBC*_) along the contralateral excitatory pathway (Figure 2A). The ITD tuning curve shifted towards the center (0-ITD) when SBC myelination increased (Figure 2B,C). The corresponding best ITD, the FWHM, and the peak firing rate of the ITD tuning curves for the left MSO were quantified and are shown in Figures 2D-E.

**Figure 2.**
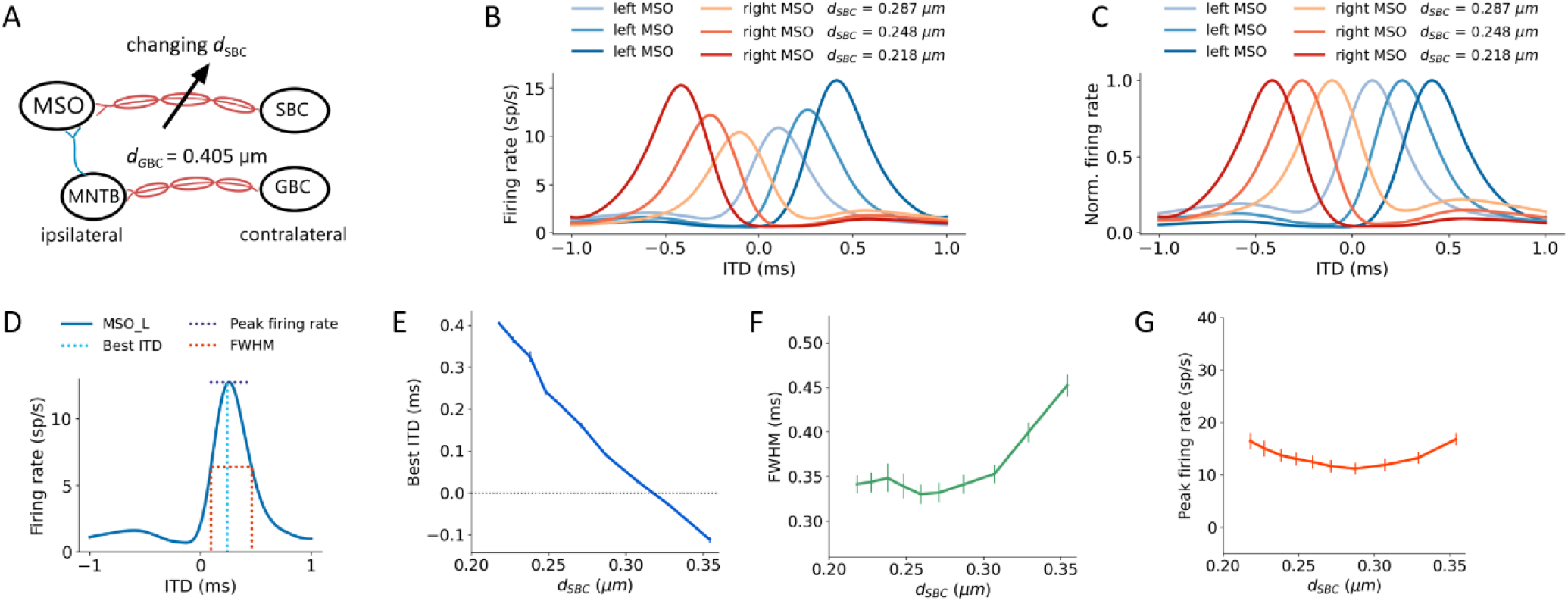
Axon myelination along the contralateral excitatory pathway and ITD tuning. (A) Sketch describing tuned parameter. (B, C) ITD tuning curve (B) and normalized ITD tuning curve (C) as a function of different SBC axon myelin thicknesses. (D) Annotation of metrics. (E) Best ITD versus SBC axon myelin thickness. (F) Full width at half maximum (FWHM) versus SBC axon myelin thickness. (G) Peak firing rate versus SBC axon myelin thickness. Error bars represent standard errors.

The best ITD decreased linearly proportional to the myelin thickness of the SBC axon (Figure 2E). A positive best ITD value indicates that the left MSO fired more rapidly when the sound stimulus first arrived at the right ear. This result is in line with experimental observations and has been described by the opponent-channel coding model (Encke and Hemmert, 2018; Magezi and Krumbholz, 2010). In addition, as the myelin thickness increased beyond 0.32 μm, the best ITD value assumed values smaller than 0 ms. This shift caused the opponent-channel coding to fail and ITD responses as well as the coding scheme to reverse, predicting a sound location on the ipsilateral side of the brainstem. This result has not been experimentally reported. The SBC myelin thickness also altered the peak width of the ITD tuning curve (Figure 2F), especially when the thickness was greater than 0.3 μm. On the other hand, ITD peak firing rates were altered by SBC myelin thicknesses on a relatively small scale (Figure 2G), where the peak firing rate dropped only about five spikes per second when the SBC myelin thickness increased from 0.22 μm to 0.3 μm.

#### GBC axon myelin thickness and ITD tunning

An increase in myelination (*d*_*GBC*_) of the GBC axon along the contralateral inhibitory pathway (Figure 3A) showed significant changes to both the shapes and the scales of the ITD tuning curves (Figure 3B,C). The best ITD value varied non-monotonically with GBC myelin thickness (Figure 3D). The best ITD value first shifted away towards the center (0 ITD), and later away from the center with a swing of approximately 100 μs. The best ITD value reached its minimum when the GBC myelin thickness was around 0.24 μm and plateaued with a higher best ITD when the thickness was thicker than 0.3 μm. The peak width decreased monotonically from 500 μs to 320 μs when the GBC myelin thickness was increased from 0.2 to 0.33 μm, and subsequently became steady with thicker GBC myelination (Figure 3E). Note that a wider FWHM indicates broader tuned ITDs for the MSO, and a narrower FWHM suggests the tuning curve has a higher sensitivity and precision. Therefore, the results from this simulation indicate that GBC axons with relatively thicker myelination can yield more precise ITDs. Finally, the peak firing rate dropped substantially from around 40 to 10 spikes per second when the GBC myelin thickness increased from 0.22 to 0.27 μm, resulting in roughly stabilized peak firing rates (Figure 3F).

**Figure 3.**
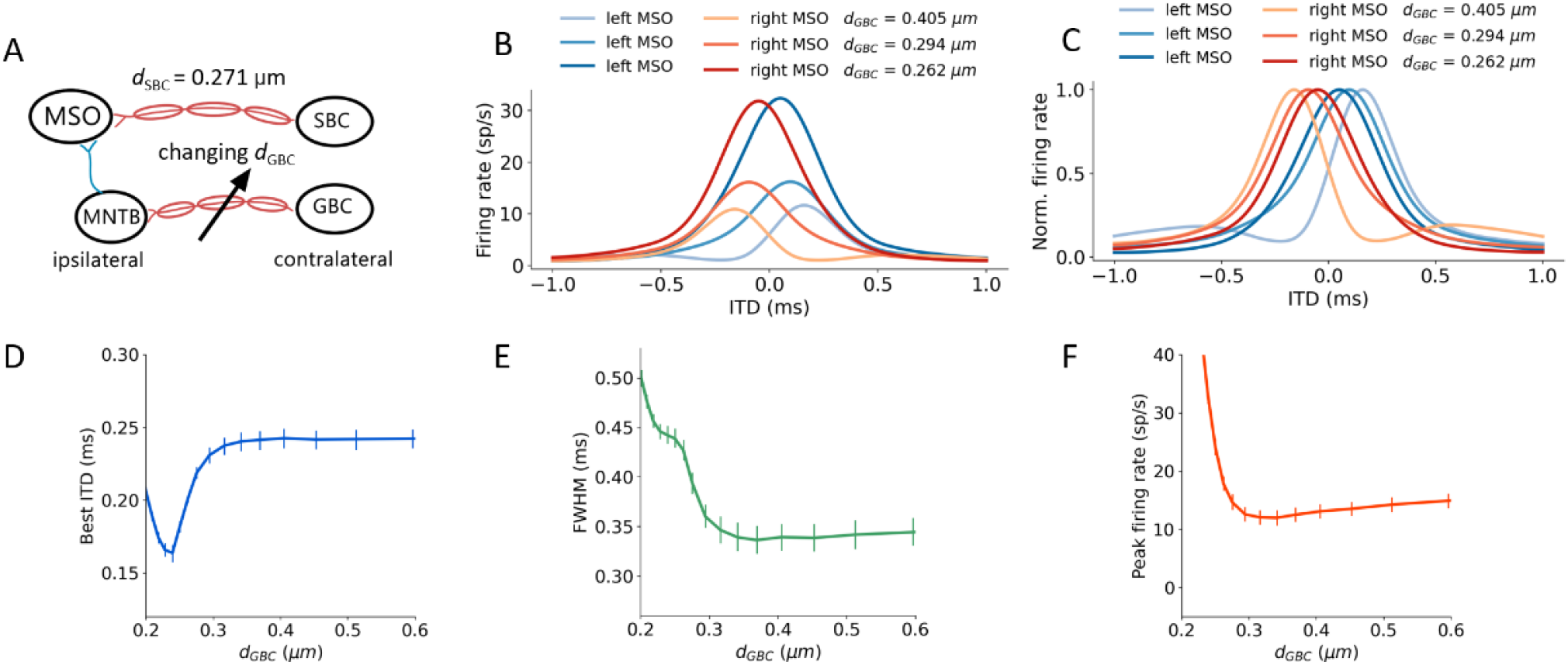
Axon myelination along contralateral inhibitory pathway and ITD tuning. (A) Sketch describing tuned parameter. (B, C) ITD tuning curve (B) and normalized ITD tuning curve (C) as a function of different GBC axon myelin thicknesses. (D) Best ITD versus GBC axon myelin thickness. (E) Full width at half maximum (FWHM) versus GBC axon myelin thickness. (F) Peak firing rate versus GBC axon myelin thickness. Error bars represent standard errors.

### 3.3 Both SBC and GBC myelination influences ITD

The interaction between myelin thickness of two contralateral inputs (one excitatory and one inhibitory) towards shaping ITD are shown in Figure 4. The increase of the SBC myelin thickness (*d*_*SBC*_) had a much more significant effect on the best ITD, where the best ITD was increasing with less SBC myelination. Similarly, GBC myelin thickness (*d*_*GBC*_) also shifted the best ITD within a specific range, especially for *d*_*GBC*_ from 0.2 to 0.3 μm (Figure 4A-C). The FWHM and peak firing rate were affected by both SBC and GBC myelin thickness. Thinner SBC myelin and thicker GBC myelin tended to produce narrower FWHM (Figure 4D-F) and lower peak firing rates (Figure 4G-I).

**Figure 4.**
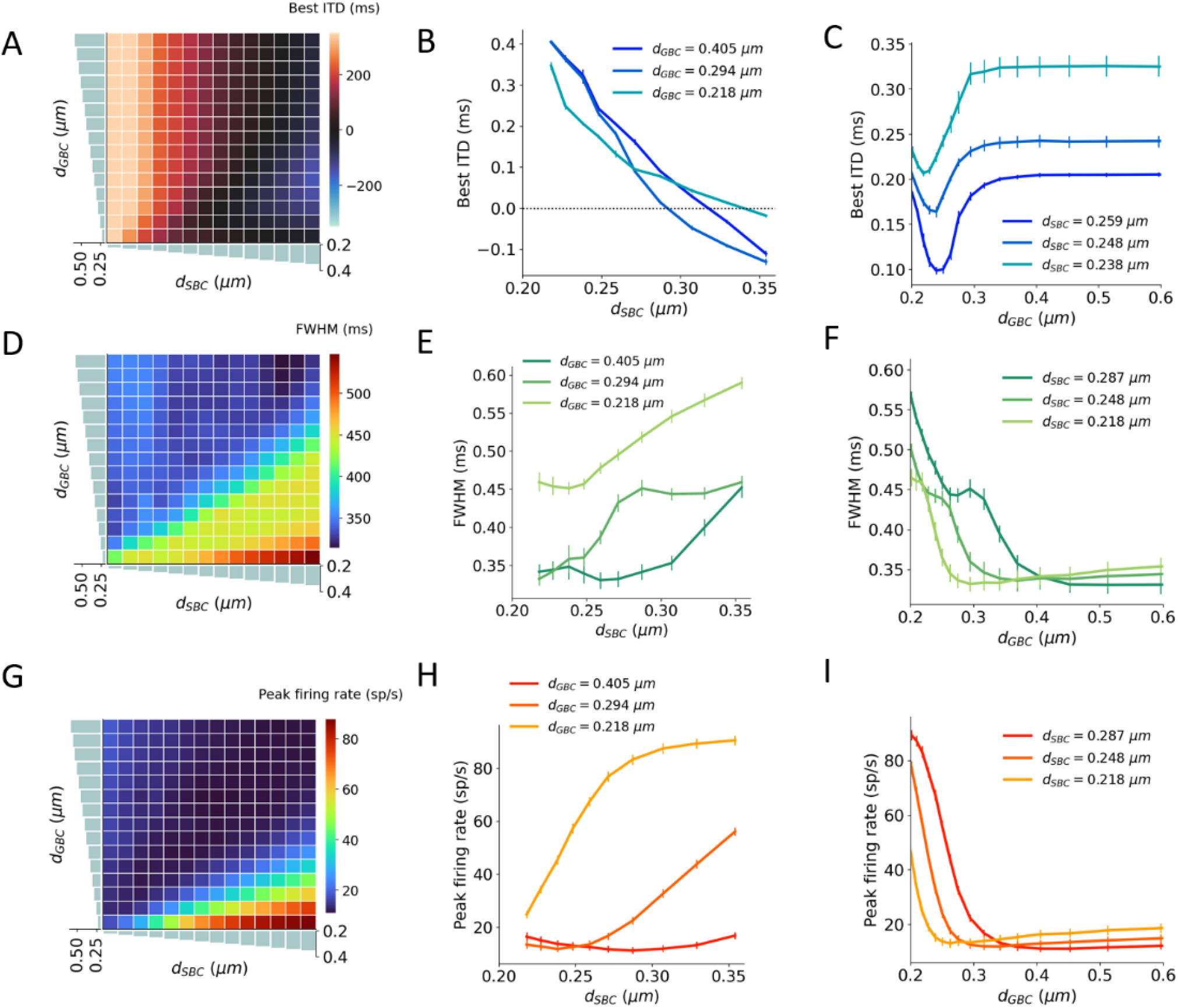
Interaction between SBC and GBC axon myelination on ITD tuning. (A, B, C) Best ITD as a function of different SBC and GBC myelin thicknesses. (D, E, F) FWHM as a function of different SBC and GBC myelin thicknesses. (G, H, I) Peak firing rate as a function of different SBC and GBC myelin thicknesses. Error bars represent standard errors.

### 3.4 Myelination affecting ITD encoding accuracy

An ITD encoding analysis was conducted using MSO neural responses to repeated pure tone stimuli with 17 different ITD steps varying from –1 ms to 1 ms. The encoding accuracy can be interpreted as the amount of ITD information extracted by the MSO (Figure 5A-C). The results indicate that the encoding accuracy reached its optimum when either SBC myelin was significantly thicker than GBC myelin, or when GBC myelin became much thicker than SBC myelin. The ability to accurately encode ITDs can also be represented as the mean squared error (MSE) between true ITDs and predicted ITDs (Figure 5D-F). Similar to the conclusion on ITD decoding accuracy, the contrast between SBC and GBC myelin thickness could result in smaller MSEs.

**Figure 5.**
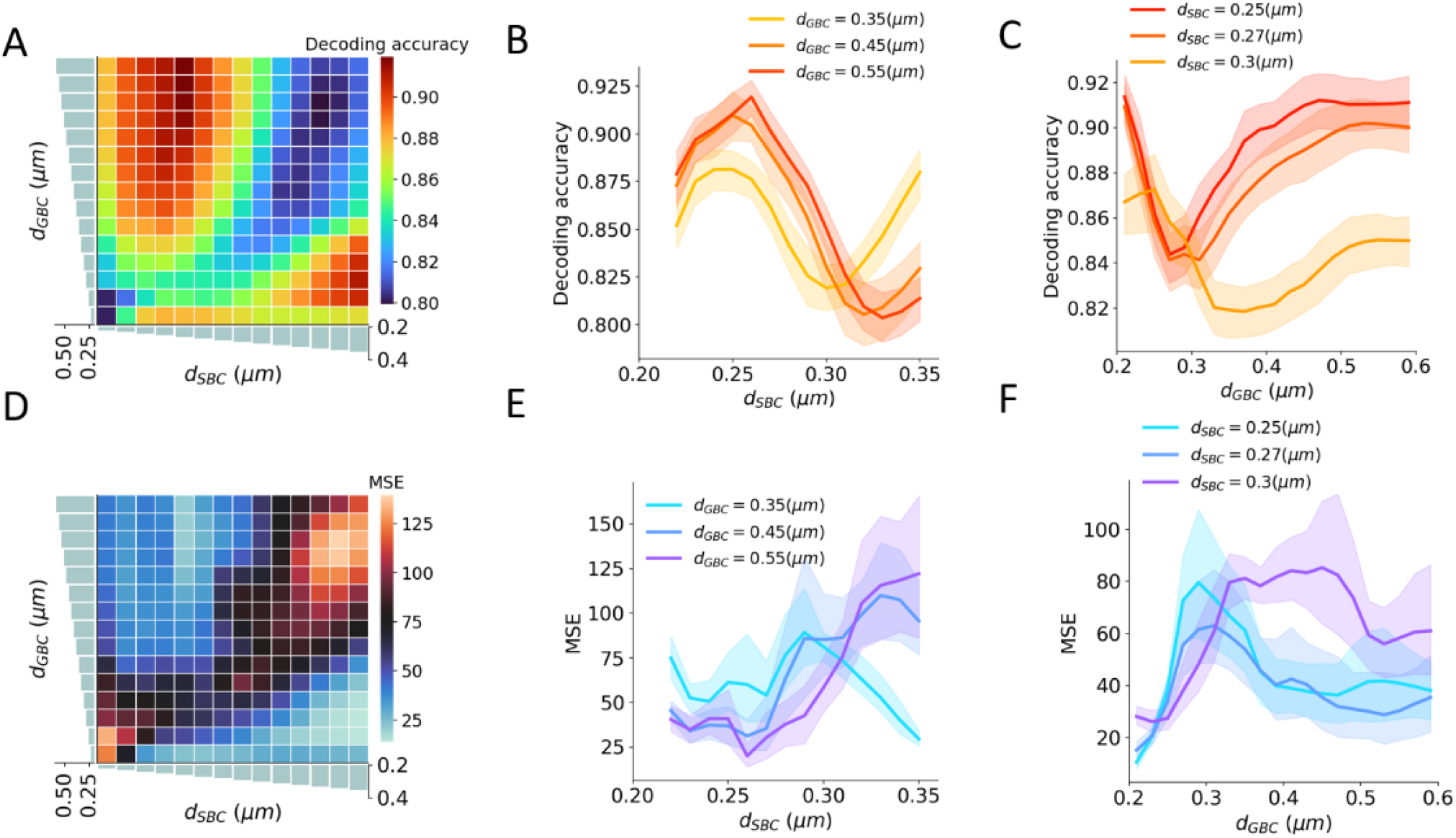
Interaction between SBC and GBC axon myelination on ITD decoding. (A, B, C) Decoding accuracy under different SBC and GBC myelin thicknesses. (D, E, F) Mean squared error (MSE) under different SBC and GBC myelin thicknesses. Error bands represent standard errors.

### 3.5 Myelin thickness and the precision of ITD perception

Natural sound clips presented with small ITDs were utilized to stimulate the circuit and calculate the precision of ITD perception. The just noticeable difference (JND) of ITD was calculated by comparing MSO responses to symmetrical ITDs, and the calculated JND can be regarded as the sensitivity to ITD stimuli. The best sensitivity was obtained when SBC myelin was thin, or GBC myelin was thick and at the same time SBC myelin was thinner than 0.3 μm (Figure 6A-C). Apart from this, the sensitivity became far worse when the best ITD approached zero. The worst sensitivity was obtained when the best ITD was negative. This result can be explained since the opponent-channel coding scheme was used in the JND calculation together with a one-tailed test. Besides calculating JND, the decoding accuracy in the range between 10 μs and –10 μs ITDs was computed to quantify the precision of the circuit to capture very small ITDs. The most precise accuracy was also acquired when the GBC myelin was much thicker than the SBC myelin (Figure 6D-F).

**Figure 6.**
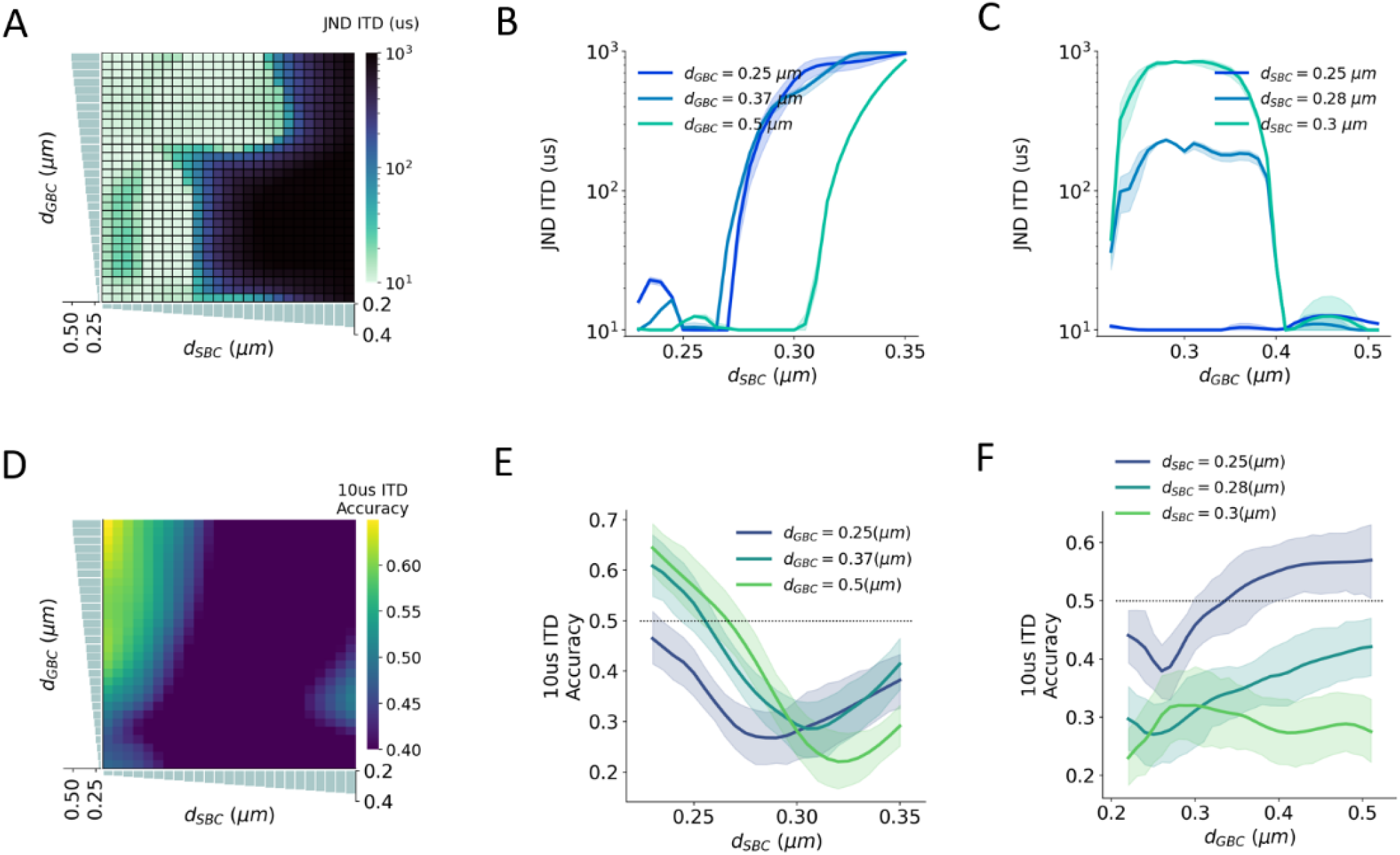
Interaction between SBC and GBC axon myelination on ITD sensitivity. (A, B, C) Just noticeable difference (JND) of ITD for different SBC and GBC myelin thicknesses. (D, E, F) Decoding accuracy of 10-us ITD for different SBC and GBC myelin thicknesses. Error bands represent standard errors.

### 3.6 The effects of contralateral inhibition on ITD perception

The SNN circuit was simulated with different synaptic strengths of the contralateral inhibitory inputs to estimate the effects of the contralateral inhibition on the ITD perception (Figure 7A). The shape of the ITD tuning curve changed as the synaptic conductance (Δ*g*_*i*_) of the MNTB to MSO increased (Figure 7B, C). The best ITD shifted closer to zero-ITD when the contralateral inhibition increased and reached a plateau with a Δ*g*_*i*_ above 25~50 nS (Figure 7D). Both peak width and peak firing rate dropped with an increasing Δ*g*_*i*_ (Figure 7E,F). Combining the results with those obtained from varying inhibition strength and myelin thickness, the maximum decoding accuracy, the minimum MSE and the minimum JND could be obtained when the SSN circuit has an optimal *ginhi* of about 50 nS and the GBC myelin was 0.2 μm thicker than the SBC myelin (Figure 7.G-I).

**Figure 7.**
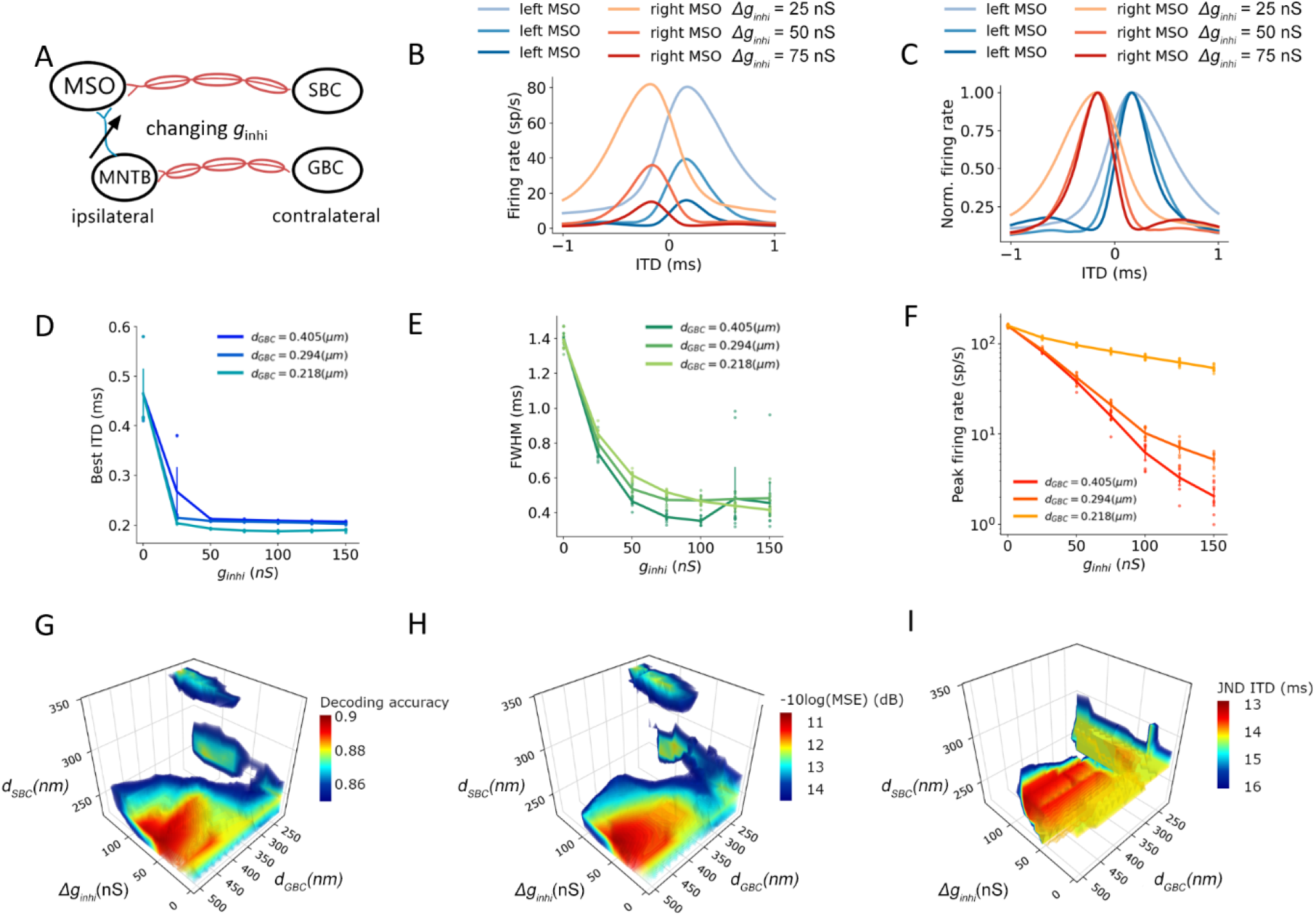
Effects of contralateral inhibition on ITD tuning and perception. (A) Sketch describing tuned parameter. (B, C) ITD tuning curve (B) and normalized ITD tuning curve (C) as a function of different contralateral inhibitory synaptic strength. (D) Best ITD versus contralateral inhibitory synaptic strength with a SBC myelin thickness of 0.248 um. (E) Full width at half maximum (FWHM) versus contralateral inhibitory synaptic strength with the SBC myelin thickness of 0.248 um. (F) Peak firing rate versus contralateral inhibitory synaptic strength with the SBC myelin thickness of 0.248 um. (G, H, I) Decoding accuracy (G), mean square error in decibel (H), and just noticeable difference of ITD (I) change with contralateral inhibitory synaptic strength and contralateral myelin thickness; values above 75% quantile were colored.

## 4 Discussion and Conclusion

### 4.1 Myelination, conduction velocity, and spike timing

In this study, the MSO circuit was modeled and tested with varying axon myelination properties to determine which role myelination plays in conduction velocity and transmission delay of action potentials. We found that changes in axon myelin thickness affected the average propagation delays of action potentials between nuclei. A thicker myelin layer on GBC axons resulted in faster action potential propagation for the contralateral inhibition, and at the same time, a thinner myelin layer on the SBC axons resulted in longer delays for the contralateral excitation to MSO.

Our decoding results (Figure 5 and Figure 6) suggest that a larger GBC myelin layer combined with a smaller SBC myelin layer produced an optimal decoding accuracy and resulted in optimal ITD precision. In this scenario, action potentials associated with contralateral inhibition propagated faster than those associated with the excitation. This phenomenon had also been reported in previous experimental studies (Brand et al., 2002; Robert et al., 2013). Specifically, it was postulated that MSO receives contralateral inhibition earlier than contralateral excitation, despite an additional synapse in the inhibitory pathway and a longer distance to cover. These findings may indicate that this tuning of myelination of axons on the contralateral pathways could be the consequence of structural adaptation of the sound localization pathway towards more accurate ITD detection.

### 4.2 Negative best ITD

The optimal decoding accuracy and small MSE can mathematically be also achieved with a completely opposite myelination pattern consisting of thicker SBC myelin and thinner GBC myelin (Figure 5). Under this opposite scheme, the peak firing rate became much higher, and the best ITDs were negatively shifted away from the zero-ITD (Figure 4). From a purely mathematical perspective, the higher peak firing rate achieved in this scheme increases the signal dynamic range, thus improving the signal-to-noise ratio for the encoded ITDs. The sign of the ITD can be thought of being irrelevant for the total amount of information, since negative or the positive best ITDs encode the same ITD value. However, from an experimental standpoint, such a non-opponent-channel coding scheme has not been described.

### 4.3 Possible roles of inhibition in MSO

Several competing models have been suggested for the role of inhibition in the MSO in ITD tuning (Grothe et al., 2010; Brande et al., 2002; Pecka et al., 2002; Couchman 2010, van der Heijden et al. 2013, Roberts et al., 2013; Myoga et al., 2014; Franken et al., 2015). Some studies support the hypothesis that this inhibition modulates the peak timing of the excitation and tunes the coincidence detection. Notably, the pharmacological blockage of inhibition shifts the best ITD towards the zero-ITD (Brand et al., 2002; Pecka et al., 2008). Moreover, conductance clamp recordings (Myoga et al., 2014) demonstrated that precisely timed inhibition could tune the best ITD by modulating the net excitatory post-synaptic potential (EPSP), and the leading contralateral inhibition biased the coincidence detection timing about 50~150 μs. The experimental result is comparable to our computational results (Figure 3D and Figure 4A,C) in which the increased GBC myelin thickness modestly shifted the best ITD by only 100 μs but resulted in much improved sensitivity and decoding accuracy.

On the other hand, other studies have challenged this inhibition-tuning model. Although well-timed leading contralateral inhibition was observed in an in-vitro study (Roberts et al., 2013), the ITD function and EPSP did not significantly differ with an inhibitory conductance of a 300-μs leading contralateral inhibition. In addition, although shift of best ITD towards zero was shown for pharmacological blockage, Roberts et al (2013) described the role of inhibition as transient and less significant over time, inferring that the removal of inhibition should not systemically shift the best ITD (Franken et al., 2015). In these studies, the occurrence of inhibition decreased overall firing rates across ITDs and narrowed the ITD functions without shifting the best ITD. This trend is consistent with our results to some degree (Figure 6), as the best ITD was not shifted with varying inhibitory synaptic conductances unless the inhibitory conductance was lower than 50 nS (Figure 6D). The increased inhibition also reduced the peak firing rate (Figure 6E) and peak width (Figure 6F) in a way similar to the increased myelin thickness and conduction velocity on the contralateral inhibitory pathway (Figure 3D-F). On the other hand, we note that the ITDs produced by varying myelination in the afferent excitatory pathway exceed the biologically relevant range of at least most mammalian species (Figure 2B,C), while the smaller range of ITDs produced by varying myelination in the inhibitory pathway matches that range closer. It is, therefore, possible that this study underestimates that role.

The relative timing of the binaural excitation was concluded to be the dominant factor for ITD tuning due to its apparent capability to regulate the best ITD compared to the inhibition (van der Heijden et al., 2013; Roberts et al., 2013; Seidl and Rubel, 2015). However, through the decoding analysis, the linkage between the best ITD and the estimated ITD sensitivity was unexpectedly shown to be more indirect but in a profound way in which the best ITD shifts could not simply be used to predict the precision of the ITD detection. Even though the presence of the leading contralateral inhibition reduced the peak firing rate and narrowed the peak width of the ITD tuning curves, the decoding results revealed that the timing and synaptic strength of the contralateral inhibition largely attributed to the pinpoint precision and sensitivity of the ITD perception (Figure 6G-I). Therefore, our findings imply that the complexity of ITD tuning depends on the temporal interaction between the excitation and inhibition.

### 4.4 Limitations

Our computational model was designed to probe the influence of axon myelination. We simplified the model and omitted several possible mechanisms of ITD encoding. First, a low-threshold potassium current shown to interact with the synaptic inhibition in MSO and sharpen the temporal sensitivity of the binaural integration (Khurana et al., 2011; Roberts et al., 2013; Myoga et al., 2014) was omitted. Furthermore, post-inhibitory facilitation that can raise the firing rate under certain conditions (Beiderbeck et al., 2018; Ma et al., 2021) and had been observed in the MSO of juvenile mice (Dodla et al., 2006) was also not considered. This phenomenon could possibly compensate for a decreased firing rate induced by the leading contralateral inhibition in MSO.

### 4.5 Conclusion

By using an SNN model of the auditory brainstem, we found that axon myelination regulated ITD perception. The myelination of contralateral excitatory pathways shifted the best ITD. Moreover, the myelination and synaptic strength of contralateral inhibition influenced the peak firing rate and width of the ITD tuning curve, and subsequently modulated the ITD precision and sensitivity.

## Acknowledgement

Supported by NIH R01 DC 17924 and R01 DC 18401. Additionally, this work utilized the Summit supercomputer, which is supported by the National Science Foundation (awards ACI-1532235 and ACI-1532236) to the University of Colorado Boulder and Colorado State University.

